# Viral Nucleosome-Like Particles Exhibit Dynamic Flexibility and Reduced Thermodynamic Stability

**DOI:** 10.1101/2025.07.22.666224

**Authors:** Melanie Melo, Jeff Wereszczynski

## Abstract

DNA packaging imposes fundamental physical constraints on genomes across the tree of life. However, most of our mechanistic understanding of these processes comes from the eukaryotic nucleosome, where highly basic histones, along with their flexible tails, coordinate DNA compaction and gene accessibility. Large DNA viruses challenge this paradigm by assembling nucleosome-like particles with a divergent histone architecture. These viral assemblies lack canonical histone tails, contain covalently fused domains linked by structured connectors, and exhibit altered surface electrostatics, features that collectively impose unique biophysical properties on viral chromatin. Here, we use multi-microsecond all-atom molecular dynamics simulations to dissect how histone fusion, tail loss, and connector architecture reshape the structural and thermodynamic behavior of the Melbournevirus nucleosome, a model system for studying viral chromatin organization. We find that viral systems exhibit elevated DNA unwrapping, weaker and more transient histone–DNA contacts, and localized flexibility at connector regions. Conformational adaptation at histone junctions partially offsets these effects, with structural shifts tuned to local DNA geometry during wrapping transitions. By capturing how nucleosome dynamics shift across time and sequence, our study reveals how conserved biophysical principles of chromatin architecture are reconfigured across life to meet distinct evolutionary demands.

## Introduction

Packaging long genomes into tight cellular spaces while maintaining regulated gene expression is a central biophysical challenge across all domains of life. In eukaryotes, this is accomplished through the hierarchical organization of chromatin,^1–3^ whose basic repeating unit is the nucleosome:^4–6^ a complex composed of approximately equal parts DNA and histone proteins. For many years, histones and nucleosomes were thought to be exclusive to eukaryotes, reflecting a specialized solution to genome organization. This view has shifted with the discovery of histone-based systems in archaea ^7–9^ and large DNA viruses,^10–15^ as well as bacterial histone-fold proteins that compact DNA through distinct, non-nucleosomal mechanisms. These findings reveal a broader principle: histone-based DNA organization is not an evolutionary outlier but a recurring solution to the mechanical and regulatory demands of genome packaging.^8,9,13,15–17^

In eukaryotes, the nucleosome is the fundamental unit of chromatin and is formed by wrapping *∼*147 base pairs of DNA around a histone octamer.^18–20^ This octamer comprises a central (H3–H4)_2_ tetramer flanked by two H2A–H2B dimers. Each histone features a conserved histone fold domain that facilitates protein–protein interactions and anchors DNA via hydrogen bonding, electrostatics, and shape complementarity. Four-helix bundle interfaces between dimers and tetramers stabilize the octamer while allowing dynamic rearrangements. Flexible N- and C-terminal tails extend from the core and mediate chromatin compaction, inter-nucleosome interactions, and recruitment of regulatory factors.^21–23^ This architecture preserves genome integrity and remains responsive to signals that modulate gene accessibility.^1^

Recent discoveries have shown that large double-stranded DNA viruses also encode histone homologs that form nucleosome-like particles.^10,24–26^ Notably, members of the Marseilleviridae family,^14^ including Melbournevirus, express fused histone doublets where analogs of the H4–H3 and H2B–H2A domains are covalently linked via short, structured connectors. These doublets retain the canonical histone fold and assemble into a nucleosome-like structure that wraps *∼*121 base pairs of DNA, forming a compact unit without linker DNA.^24,25,27^ Although the overall nucleosome architecture is preserved, including the tetramer–dimer interface and four-helix bundles, histone fusion may restrict local flexibility within each histone pair, and the structured connectors likely impose new dynamic constraints at their junctions. Many viral histones also lack the long, disordered terminal tails found in eukaryotic counterparts, eliminating key contacts that regulate higher-order chromatin organization and modulate DNA accessibility. Together, these features define a minimalist but physically distinct chromatin architecture^15^ that challenges conventional models of nucleosome behavior and raises critical questions about how dynamic instability and structural plasticity are encoded in tail-less, fused histones.

Despite recent structural insights from cryo-EM,^24,25,27^ the dynamic behavior of viral nucleosomes remains largely uncharacterized. Observations of partially unwrapped DNA, histone fusions, and the absence of terminal tails all point to a chromatin architecture that may behave quite differently from its eukaryotic counterpart. Electrostatic surface calculations highlight this difference: viral histones exhibit a markedly more neutral charge distribution than eukaryotic histones, consistent with weaker intrinsic DNA affinity (Figure 1). This shift in electrostatics could alter the balance of forces that govern nucleosome unwrapping,^28,29^ compaction, and conformational plasticity.^30^ Moreover, the structured connectors linking histone domains in viral systems may restrict or redirect local flexibility in ways that remain undefined.

**Figure 1:**
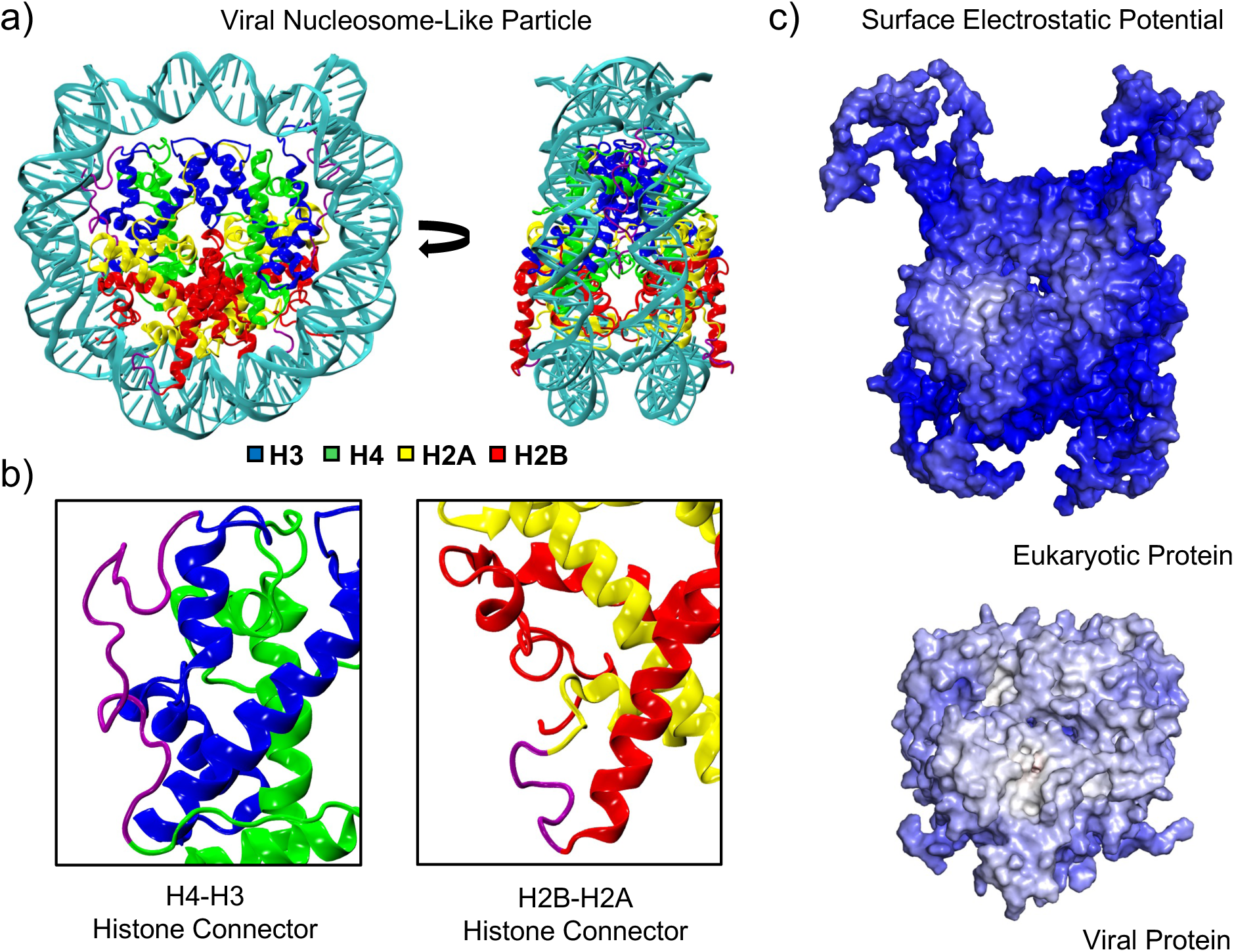
**(a) Melbournevirus nucleosome-like particle structure.** The overall structure features fused histone doublets wrapping DNA into a compact, nucleosome-like particle. **(b) Histone connector architecture.** Close-up of histone connectors covalently linking H4–H3 and H2B–H2A domains, forming fused histone doublets. **(c) Electrostatic surface potentials.** Eukaryotic and viral histone cores were calculated using the Adaptive Poisson–Boltzmann Solver (APBS).^38^ The eukaryotic core exhibits a strongly basic outer surface where acidic DNA binds, stabilizing the nucleosome. In contrast, the viral core is more electrostatically neutral in this region, weakening the protein–DNA interface. Potentials are plotted on a red–white–blue scale from –30 to +30 kT/e.

While current structural models have revealed key architectural details, they cannot capture how these assemblies behave over time or respond to sequence-specific and mechanical cues. A dynamic perspective is essential for uncovering how physical features like DNA unwrapping, histone flexibility, and connector mobility shape chromatin behavior.^27,30^ Here, we use multi-microsecond all-atom molecular dynamics (MD) simulations to dissect the dynamic properties of viral chromatin. Such simulations have yielded critical insights into eukaryotic nucleosome behavior, including DNA breathing, histone variant incorporation, tail dynamics, and unwrapping transitions.^31–37^ By comparing nucleosomes containing canonical histones, truncated eukaryotic histones, and fused viral histone doublets, we isolate the physical consequences of tail loss, histone fusion, and connector architecture. Although often referred to as nucleosome-like particles, we use the term “viral nucleosomes” for brevity. Likewise, while the Melbournevirus fused viral histones are typically labeled Ha–HA and Hv–Ha, we adopt the eukaryote-based H4–H3 and H2B–H2A notation to reflect structural homology (Figure S1). We find that viral nucleosomes exhibit substantially more DNA unwrapping, weaker and less persistent histone–DNA contacts, and greater flexibility in connector regions. These differences shape distinct unwrapping pathways and indicate that the viral nucleosome’s energetic landscape is shifted to favor local plasticity over global stability, diverging from canonical chromatin behavior. Our findings clarify how viral chromatin achieves tight genome packing^15^ while retaining the capacity for regulated access and expand the physical principles that govern genome organization across evolutionary lineages.

## Methods

### System Preparation

Three nucleosome systems were constructed for molecular dynamics (MD) simulations: a full-length eukaryotic nucleosome, a truncated eukaryotic nucleosome, and a viral nucleosome-like particle. Each nucleosome type was constructed with both the Widom 601,^39^ and Alpha-Satellite DNA sequences,^40^ yielding six distinct systems in total.

The full-length eukaryotic model was based on PDB 1KX5,^19^ which contains all four histone proteins with intact N- and C-terminal tails and 147 base pairs of Alpha-Satellite DNA. The truncated eukaryotic system was generated by deleting the disordered N- and C-terminal tail regions of each histone: 42 residues from H3, 28 from H4, 43 from H2A, and 32 from H2B. Truncations were applied to the N-termini of all histones, and to the C-terminus of H2A, consistent with previous tail-deletion protocols.^23^

The viral nucleosome-like particle was modeled from PDB 7N8N,^24^ a cryo-EM structure of the Melbournevirus histone complex wrapped with Widom 601 DNA. Because the deposited structure only resolves 129 of the expected 147 base pairs, DNA extensions were manually built in Visual Molecular Dynamics (VMD).^41^ To guide the system toward a closed, fully wrapped conformation, targeted MD^42^ was applied, gradually pulling the heavy atoms of the DNA residues toward a reference structure based on the 1KX5 structure with a force constant of 1.0 kcal/mol *·* Å^-2^. Both the fitting and RMSD calculation for the restraint were performed on this same atom selection, achieving a final RMSD of 0.64 Å relative to the restrained DNA atom selection. The resulting model achieved a cross-correlation coefficient of 0.62 against the experimental cryo-EM density (EMD-24238),^24^ indicating good agreement given the resolution and flexibility of the modeled DNA extensions.

All systems were constructed in tleap (AmberTools)^42^ with the OPC water model^43^ and neutralized with Na^+^ and Cl^−^ to a final ionic strength of 150 mM NaCl. The ff19SB force field^44^ was used for proteins, and BSC1 for DNA.^45^ Hydrogen mass repartitioning was applied to enable the use of a 4 fs time step.^46^

### Molecular Dynamics Simulations

All minimization, heating, equilibration, and production simulations were run using the GPU accelerated version of PMEMD in Amber 20.^42,47^ Each system was minimized in two stages: first with 5,000 steps of steepest descent followed by 5,000 steps of conjugate gradient with a 10 kcal/mol·Å^2^ harmonic restraint applied to all nucleosome heavy atoms. A second minimization was then performed without restraints using the same protocol. Systems were heated from 5K to 300K over 10ps in the NPT ensemble using Langevin dynamics with a collision frequency of 2 ps^−1^ and a Berendsen barostat,^48^ maintaining the same positional restraints. A 10 Å cutoff was used for short-range nonbonded interactions, and particle mesh Ewald was used to treat long-range electrostatics.^49^ Following heating, each system underwent multi-phase equilibration at 300 K in the NPT ensemble using a 4 fs timestep, with harmonic restraints sequentially reduced over seven equilibration phasesd: 10.0, 3.0, 1.0, 0.3, 0.1, 0.03, and 0.01 kcal/mol·Å^2^. Each phase lasted 100–200 ps, followed by a final 1 ns unrestrained equilibration.

Production simulations were run under the same NPT conditions. For the full-length eukaryotic systems, an additional 300 ns simulation was first performed to allow histone tail collapse prior to initiating production runs. Then, for all systems, four independent 2.0 µs production simulations were performed to increase reproducibility and the reliability of error estimates.^50^ The first 100 ns of each simulation were excluded from analysis to allow for equilibration. Simulations were performed using local GPU clusters, the Argonne Leadership Computing Facility (ThetaGPU), and XSEDE resources.^51^

### Trajectory Analysis

Structural and dynamic properties of each system were assessed using CPPTRAJ^52^ from AmberTools^42^ and the MDAnalysis Python package.^53^ All analyses focused on heavy atoms unless stated otherwise and excluded histone tails unless explicitly included. Visualization and figure generation were performed using VMD^41^ and Python-based plotting libraries.^54,55^

To quantify structural drift over time, trajectories were aligned to the histone core (Figure S1), and the root-mean-square deviation (RMSD) was computed. Local flexibility was assessed via root-mean-square fluctuations (RMSF) for each histone type, enabling consistent comparisons across systems and replicates. DNA unwrapping was evaluated using two complementary metrics. First, the radius of gyration (Rg) was computed separately for the Entry (base pairs 1–73) and Exit (base pairs 75–147) DNA regions. Second, a custom MDAnalysis script identified base pairs as unwrapped if displaced by more than 7 Å from their initial position and lacking contact with the histone core or adjacent gyre. Protein–DNA contacts were defined using a 4.5 Å cutoff between heavy atoms. A Savitzky–Golay filter with a first-degree polynomial and a 100 ns window was applied to smooth the unwrapping traces.

Hydrogen bond occupancies were calculated using a 3.0 Å donor–acceptor distance and a 20*^→^* angle cutoff and averaged across replicates. Connector dynamics in viral histones were analyzed via DBSCAN clustering^56^ of backbone and sidechain dihedral angles. Analyses focused on H4–H3 connectors, with trajectories aligned to the histone core and clustering performed every 1 ns. DBSCAN parameters included a minimum point density of 4 and an epsilon of 3.8. Separate analyses for Entry- and Exit-unwrapped states yielded 21 clusters in Widom DNA and 17 clusters in Alpha-Satellite DNA.

Molecular mechanics generalized Born surface area (MM/GBSA) analyses were performed using an igb value of 5^57^ and a 0.15 M salt concentration with the MMPBSA.py program^58^ in AmberTools24.^42^ The single-trajectory approach was applied, treating nucleic acid residues as the ligand and protein residues as the receptor. Timeseries data were analyzed using the pymbar Python package^59^ to estimate statistical inefficiency, resulting in a 100 ns decorrelation interval for all systems and energy terms. Reported error bars represent the standard error of the mean.

## Results

### Viral nucleosomes exhibit enhanced DNA unwrapping and flexibility

We assembled three distinct nucleosome protein cores for all-atom simulations: viral nucleosomes with fused histone doublets, eukaryotic nucleosomes with intact histone tails, and eukaryotic nucleosomes with truncated tails. All systems contained 147 base pairs of DNA,^60^ using either the Widom 601 or Alpha-Satellite sequence, yielding six total systems. For each, we ran four independent 2 µs all-atom molecular dynamics simulations. The viral nucleosomes were derived from cryo-EM structures of Melbournevirus nucleosome-like particles and featured fused H4–H3 and H2B–H2A connectors in place of canonical subunit interfaces, allowing direct comparison to eukaryotic systems under varied sequence-dependent DNA binding environments.

Viral nucleosomes showed highly dynamic, asymmetric DNA unwrapping,^28,29^ with the Entry and Exit ends behaving independently (Figure 2). In most simulations, one or both ends unwrapped by up to 23 base pairs and remained extended for up to 1800 ns. Entry DNA typically unwrapped rapidly to *∼*15 base pairs, though one replicate showed large fluctuations between 5–25 base pairs before stabilizing. Exit DNA was more variable: some trajectories maintained unwrapping of 15–20 base pairs, while others oscillated between 3–15 base pairs in patterns characteristic of DNA breathing. This asymmetric dynamic behavior was consistent across both Widom 601 and Alpha-Satellite sequences, indicating that it is a general feature of the viral nucleosome architecture.

**Figure 2:**
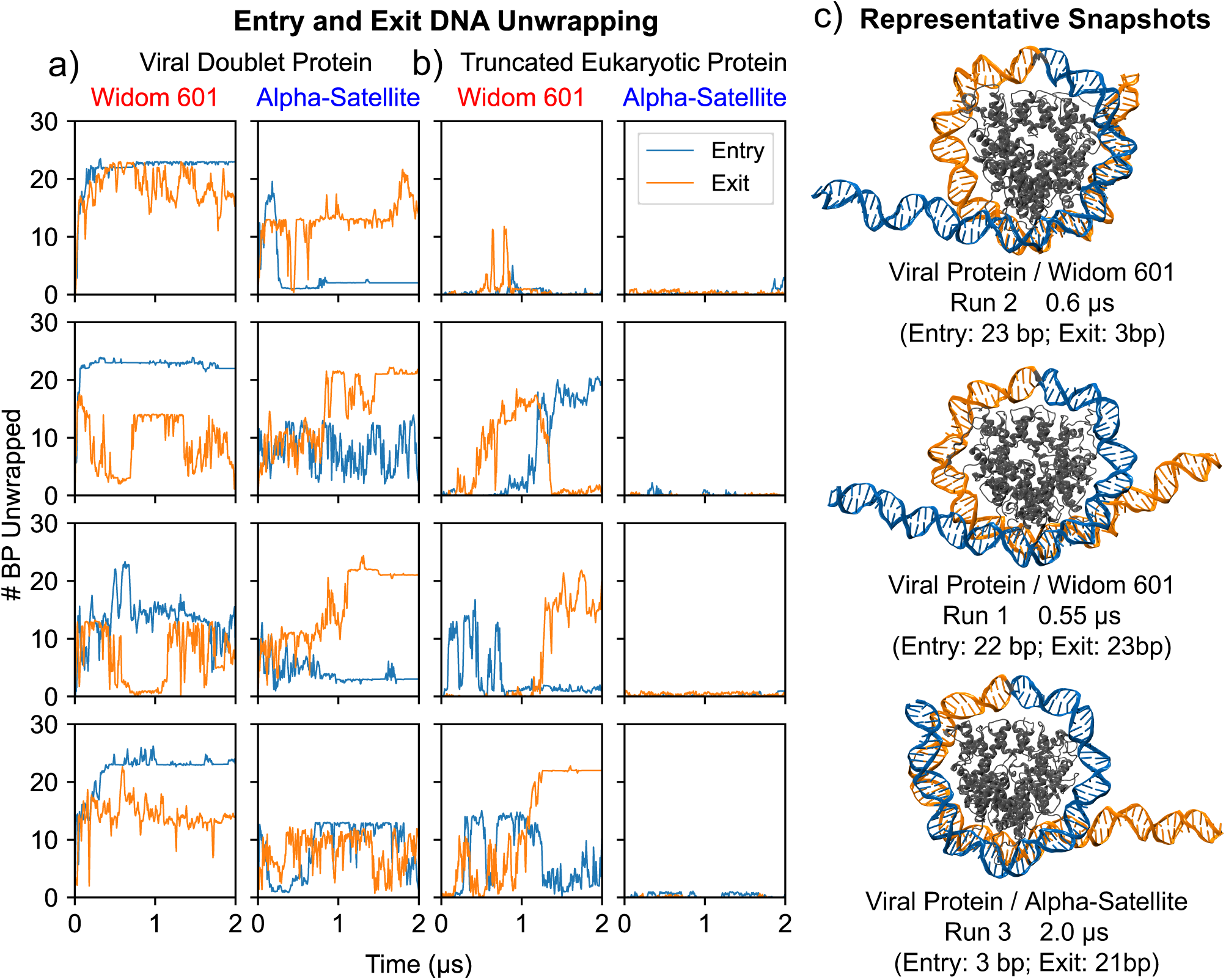
**(a) DNA unwrapping dynamics for viral histone doublets.** Widom 601 and Alpha-Satellite DNA show asymmetric unwrapping, with Entry DNA stabilizing near 23 base pairs unwrapped and Exit DNA exhibiting more dynamic behavior. **(b) DNA unwrapping dynamics for truncated eukaryotic proteins.** Alpha-Satellite DNA remains tightly wrapped (0–3 bp unwrapped), while Widom 601 shows dynamic end fluctuations up to 25 bp. **(c) Representative structural snapshots.** Snapshots show full unwrapping at the Entry site, Exit site, and both arms simultaneously.

Although total unwrapping lengths were similar between DNA sequences, the kinetics and stability of unwrapping varied substantially. Viral nucleosomes with Widom 601 DNA unwrapped rapidly at both ends, while those with Alpha-Satellite DNA exhibited slower unwrapping, typically reaching *∼*20 base pairs only after 1000–1800ns (Figure 2). In Alpha-Satellite systems, Entry DNA often stabilized near 5 base pairs or transiently extended to 15 base pairs before rewrapping. Only one replicate exhibited relatively symmetrical unwrapping, with both ends fluctuating between 3–15 base pairs throughout the simulation. These results demonstrate that viral nucleosome unwrapping is highly sequence-sensitive, with DNA composition modulating both the rate and symmetry of release.

Eukaryotic nucleosomes with full-length histones, by contrast, had minimal unwrapping across all replicates, regardless of DNA sequence. Removing the tails enabled substantial DNA release in Widom 601 systems (Figure S2). Three of four replicates alternated between fully wrapped and *∼*20–25 base pair unwrapped states at both Entry and Exit ends (Figure 2), while the fourth showed only minor, transient fluctuations. Truncated Alpha-Satellite nucleosomes remained tightly wrapped, suggesting that DNA sequence modulates unwrapping even in the absence of histone tails.

Cumulative unwrapping measurements captured the overall extent of DNA release and underscored the influence of sequence composition on unwrapping dynamics (Figure S2). Radius of gyration (Rg) analysis paralleled these findings: viral nucleosomes exhibited elevated and more variable Rg values, consistent with asymmetric unwrapping and increased sequence-dependent mobility (Figure S3). In contrast, eukaryotic systems with full-length histones maintained low, uniform Rg values (*∼*44.0 Å), reflecting persistent DNA wrapping. Together, these results underscore the enhanced conformational flexibility and sequence-sensitive dynamics that distinguish viral nucleosomes from canonical eukaryotic systems.

### Viral histone dynamics disrupt nucleosome core stability

Having established that viral nucleosomes exhibit enhanced DNA mobility, we next assessed whether this behavior was accompanied by increased flexibility within the histone core. We quantified residue-level motion using root mean square fluctuations (RMSF) of Ca atoms over the final 1.5 µs of each simulation (Figure 4). Compared to eukaryotic systems, viral nucleosomes displayed increased dynamics across the histone fold domains, particularly in regions involved in histone–DNA and histone–histone contacts. The aN helices of H3 and H2A, the L1–L2 loops, and the H2A docking domain were among the most flexible. This dynamic behavior was further modulated by DNA sequence: systems built with Widom 601 DNA consistently showed RMSF values 1–2 Å higher than those with Alpha-Satellite. Eukaryotic histones, in contrast, remained stable across sequences and truncation states, with negligible fluctuations throughout the core.

**Figure 3:**
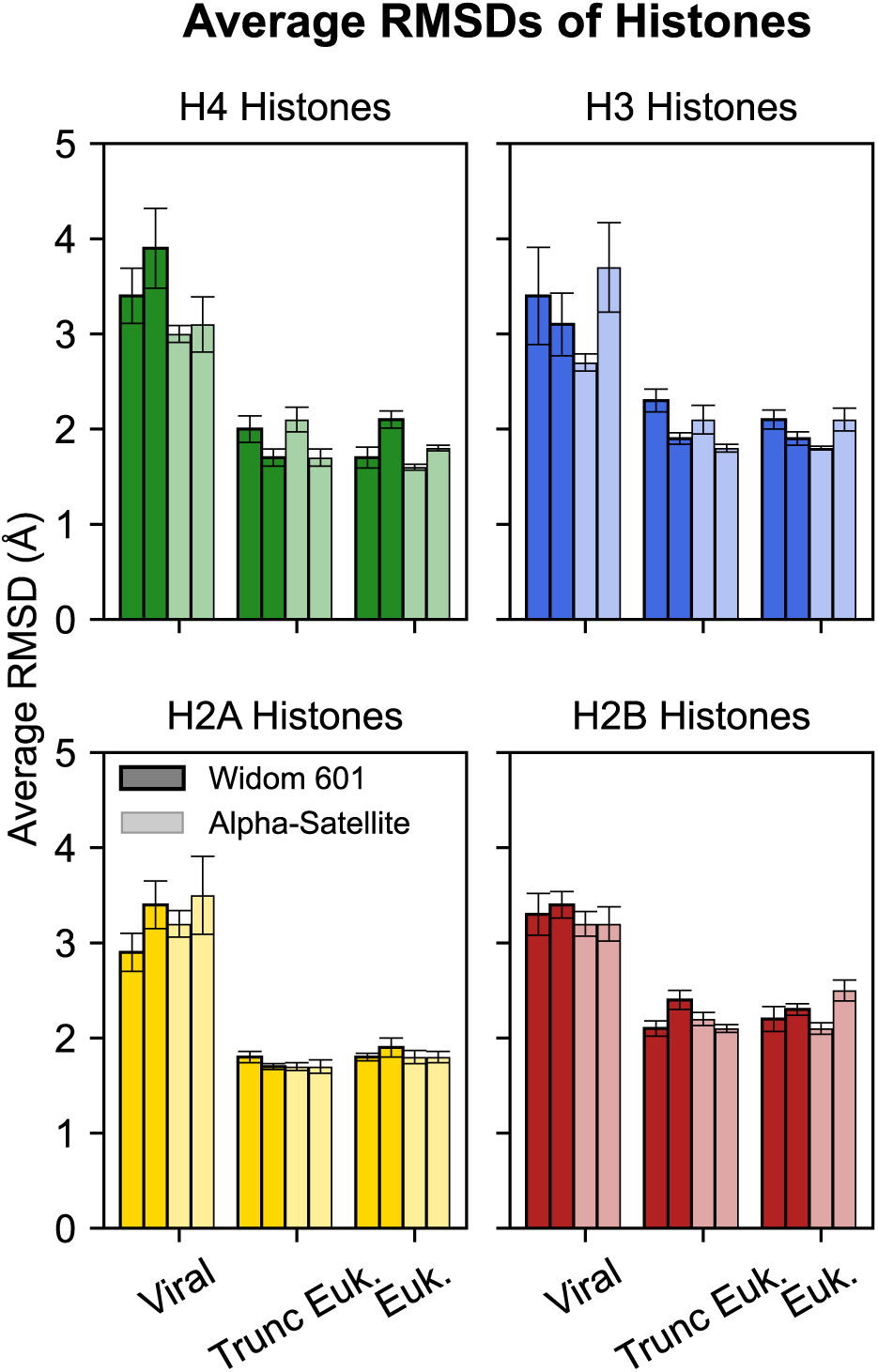
Average root-mean-square deviation (RMSD) for each histone fold domain. Viral histones show uniformly higher RMSDs than their eukaryotic counterparts, indicating greater flexibility driven by fused architecture and divergent primary sequence.

**Figure 4:**
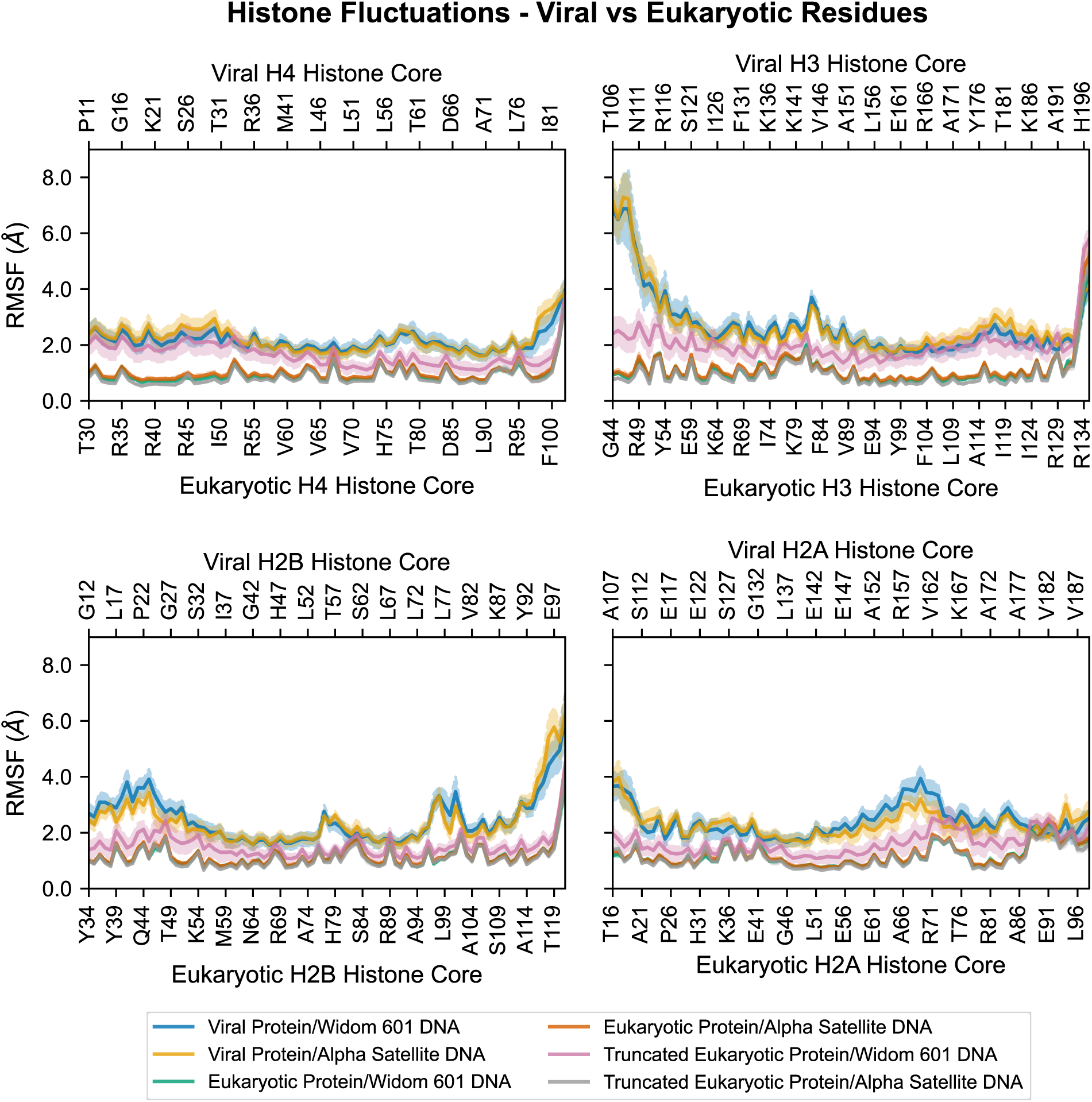
Root-mean-square fluctuation (RMSF) of histone core residues. Viral proteins show greater flexibility than eukaryotic histones, independent of tail presence. DNA sequence has minimal effect in eukaryotes but leads to variable fluctuations in viral cores.

To determine whether these local fluctuations led to broader conformational shifts, we calculated average root mean square deviations (RMSDs) of each histone fold domain during the equilibrated simulation windows (Figure 3). Viral histones consistently deviated more than their eukaryotic counterparts across all domains. H4 showed the largest average shifts (3.0–3.9 Å), followed by H2A (2.9–3.5 Å), H3 (2.7–3.7 Å), and H2B (2.6–3.1 Å). In contrast, all eukaryotic domains remained below 2.5 Å with a mean value of 1.9 Å. Truncating the eukaryotic tails resulted in mild changes to these RMSD values, all within the standard error of the mean and still well below those observed in viral systems. These differences suggest that dynamic instability in viral histones arises from their primary sequence and fused architecture, not merely the absence of flexible tails.

We then examined whether the elevated histone flexibility in viral systems translated into large-scale rearrangements of the nucleosome core. Eukaryotic assemblies, whether full-length or tail-truncated, maintained compact architectures with protein core Rg values centered near 26.5 Å (Figure 5a) and minimal relative displacements between the dimers and the tetramer. Viral nucleosomes, by contrast, exhibited broader Rg distributions with primary populations near 26.8 Å and secondary peaks above 27.5 Å in Alpha-Satellite systems, alongside increased dimer displacements of 2–4 Å (Figures S4 & S5). These shifts reflect a more flexible and heterogeneous histone arrangement that departs from the conserved eukaryotic geometry and likely facilitates the enhanced DNA unwrapping observed earlier.

**Figure 5:**
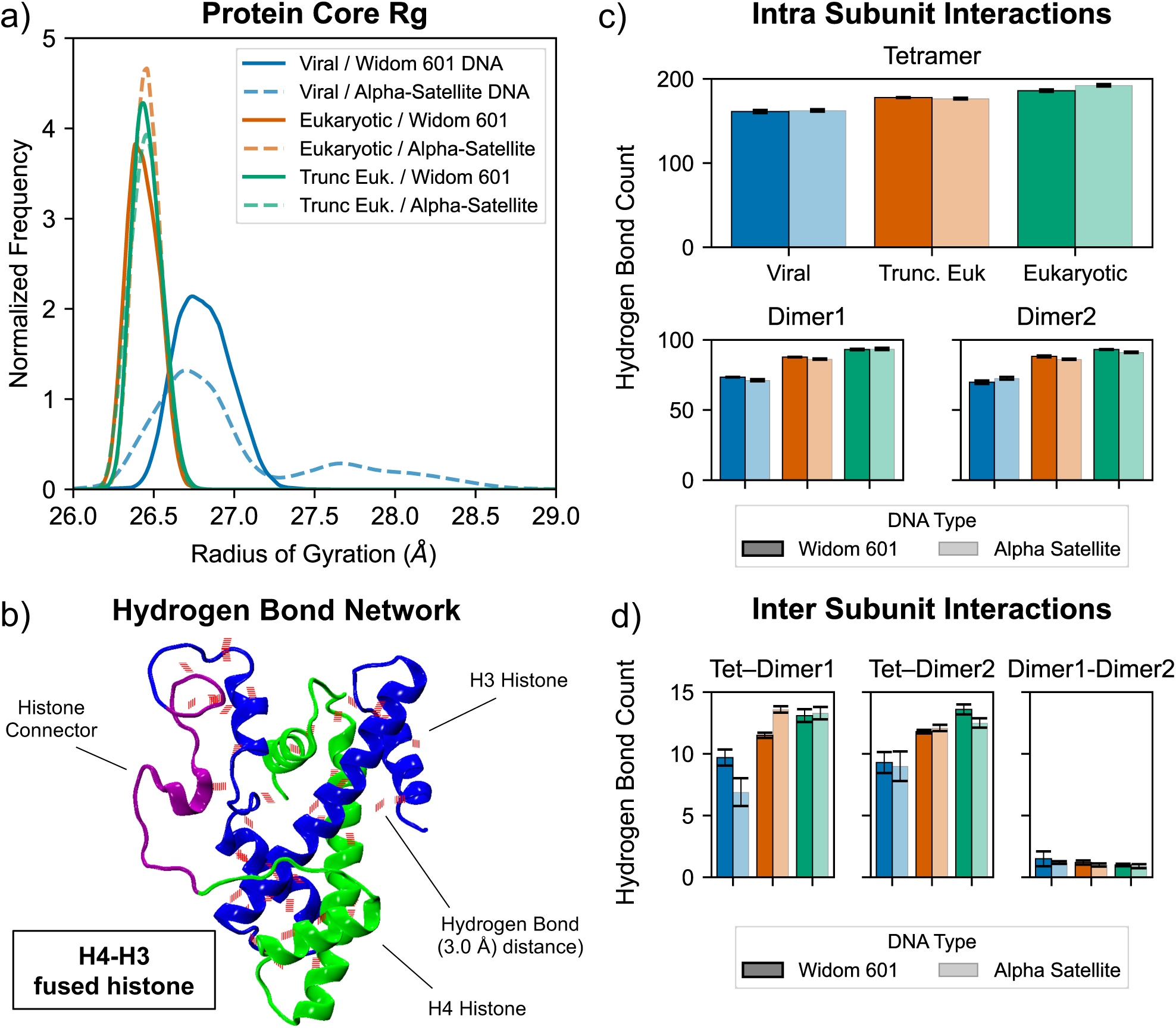
**(a) Radius of gyration (Rg) distributions.** Viral systems have reduced protein core compaction compared to eukaryotic and truncated variants. **(b) Representative hydrogen bond network.** Fused H4–H3 histones form intramolecular hydrogen bonds that contribute to structural stabilization. **(c–d) Average hydrogen bonding.** Number of hydrogen bonds within and between octamer subunits across systems (Viral, Eukaryotic, and Truncated Eukaryotic) and DNA types (Widom 601 and Alpha-Satellite).

We assessed how histone architecture influences nucleosome thermodynamics by calculating MM/GBSA binding free energies between the histone assemblies and DNA. Viral nucleosomes exhibited the weakest interactions, with binding energetics in the viral Widom 601 system of –467.1*±*23.8 kcal/mol, driven by van der Waals interactions (–567.9*±*11.4 kcal/mol) and offset by highly unfavorable electrostatics (+100.9*±*20.9 kcal/mol) (Table 1). In contrast, the eukaryotic Widom 601 system showed a threefold stronger interaction energy of –1567.9*±*24.7 kcal/mol, supported by both larger van der Waals contributions (–1502.2*±*12.0 kcal/mol) and favorable electrostatics (–65.7*±*21.6 kcal/mol). Eukaryotic systems lacking tails displayed intermediate energetics. Although MM/GBSA has multiple limitations and its absolute energies should be interpreted qualitatively, the magnitude and direction of the differences point to substantial changes in core stability (Table S1). Viral histone cores are markedly less cohesive, with weakened van der Waals contacts and repulsive electrostatics aligning with their enhanced dynamics and loosened architecture. While tails and basic residues are known to stabilize eukaryotic nucleosomes, their absence in viral systems contributes to weakened and less persistent DNA binding.

**Table 1:**
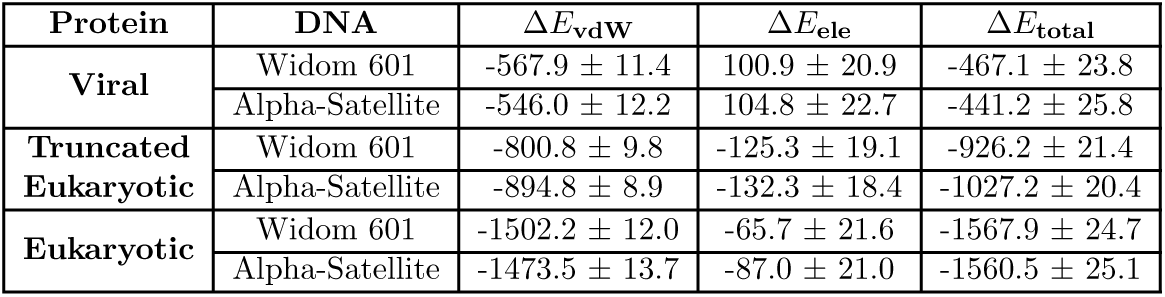
MM/GBSA binding free energies between the complete protein and the DNA interface (kcal/mol). Viral nucleosome-like particles show significantly weaker DNA binding than eukaryotic nucleosomes, driven by reduced van der Waals contributions and less favorable electrostatics.

### Altered Intra-Core Contacts Disrupt Canonical Histone Interfaces

The reduced binding energy observed in viral systems prompted us to examine the molecular interactions underlying this destabilization. Hydrogen bond and salt bridge analyses revealed systematic differences in intra-core stabilization. Viral tetramers formed fewer intratetramer hydrogen bonds—around 162 on average, compared to 189 in full-length and 177 in tailless eukaryotic systems (Figure 5c). Similar patterns were seen within dimers: viral systems averaged 72 hydrogen bonds, while full-length and tailless eukaryotic systems had 93 and 87, respectively. The reduced contact density reflects less cohesive intra-subunit packing in viral histones (Figure S6). This loosening likely underlies both their increased flexibility and their diminished thermodynamic stability.

At the tetramer–dimer interface, viral systems formed fewer and shorter-lived stabilizing interactions. Eukaryotic nucleosomes consistently formed 13 hydrogen bonds at this interface, compared to just 7–9 in viral systems, depending on sequence (Figure 5d). Additional stabilizing features were also reduced. In eukaryotic systems, the H3 a2 helix and loop 3 formed persistent salt bridges with the H2A a3 helix, including dual contacts via H3 R134 (Table 2). These high-occupancy interactions were absent in viral systems, which instead formed alternative, often weaker, contacts involving the H3 a2 helix and the H4–H2A interface. Notably, viral histones established unique hydrogen bonds between H4 loop 3 and H2A loop 3, suggesting compensatory rewiring of the interface to accommodate architectural differences.

**Table 2:**
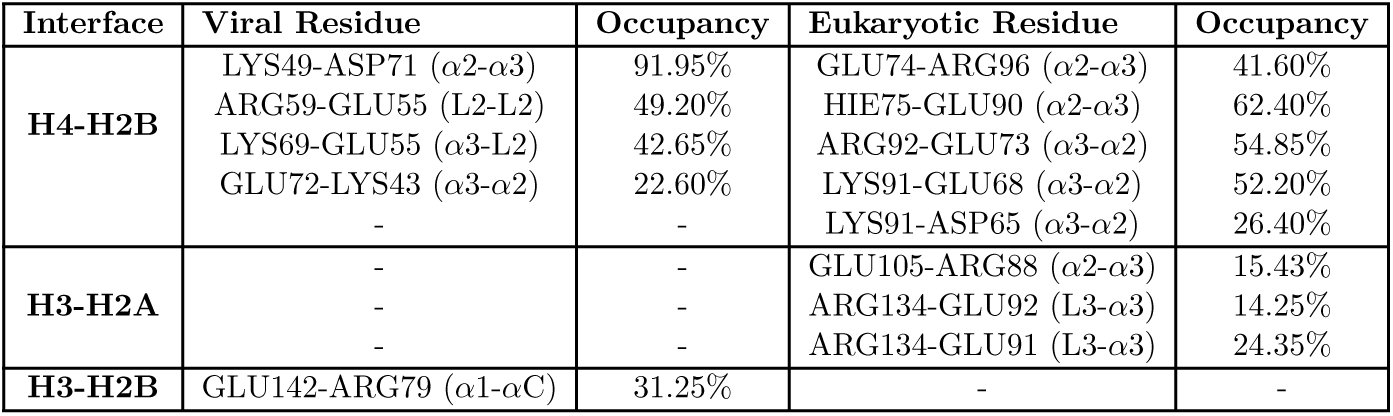
Tetramer–Dimer Interface Salt Bridges. Eukaryotic histones form stable H3–H2A salt bridges that are absent in viral systems. Viral particles instead exhibit weaker, rewired contacts—particularly at the H4–H2A and H4–H2B interfaces—reflecting reduced interface stability.

| Interface | Viral Residue | Occupancy | Eukaryotic Residue | Occupancy |
| --- | --- | --- | --- | --- |
| H4-H2B | LYS49-ASP71 ($\alpha 2$-$\alpha 3$) | 91.95% | GLU74-ARG96 ($\alpha 2$-$\alpha 3$) | 41.60% |
| | ARG59-GLU55 (L2-L2) | 49.20% | HIE75-GLU90 ($\alpha 2$-$\alpha 3$) | 62.40% |
| | LYS69-GLU55 ($\alpha 3$-L2) | 42.65% | ARG92-GLU73 ($\alpha 3$-$\alpha 2$) | 54.85% |
| | GLU72-LYS43 ($\alpha 3$-$\alpha 2$) | 22.60% | LYS91-GLU68 ($\alpha 3$-$\alpha 2$) | 52.20% |
| | - | - | LYS91-ASP65 ($\alpha 3$-$\alpha 2$) | 26.40% |
| H3-H2A | - | - | GLU105-ARG88 ($\alpha 2$-$\alpha 3$) | 15.43% |
| | - | - | ARG134-GLU92 (L3-$\alpha 3$) | 14.25% |
| | - | - | ARG134-GLU91 (L3-$\alpha 3$) | 24.35% |
| H3-H2B | GLU142-ARG79 ($\alpha 1$-$\alpha C$) | 31.25% | - | - |

Interface stabilization at the H4–H2B boundary also diverged. While four of five canonical salt bridges were conserved, viral systems showed lower occupancy overall. The viral H4 a2–H2B a3 interaction reached 92% occupancy, but eukaryotic systems displayed two moderately persistent salt bridges in the same region (42% and 62%). Viral nucleosomes also formed a noncanonical L2–L2 salt bridge between H4 and H2B, potentially contributing localized interface rigidity. At the H3–H3 four-helix bundle, eukaryotic systems formed two stabilizing contact points, while viral cores maintained only a single, transient contact—reinforcing their lower internal cohesion.

Despite comparable a-helical content and amino acid composition (e.g., *∼*50% nonpolar across homologous regions; Figure S7), viral and eukaryotic nucleosomes employ fundamentally distinct stabilization strategies. Eukaryotic cores rely on persistent hydrogen bonding among basic residues, while viral systems compensate for these interactions with transient polar and nonpolar contacts. These differences in molecular interaction networks help explain the structural heterogeneity and increased flexibility characteristic of viral histone cores.

### Viral Connector Dynamics and Energetics Reveal Limited Compensation for Histone Tail Loss

Our analyses thus far revealed that viral nucleosomes are structurally and thermodynamically less stable than their eukaryotic counterparts, with loosened histone interfaces, increased flexibility, and reduced DNA contacts. This raised the question of whether their unique architectural features help compensate for this instability. Melbournevirus systems contain covalent peptide linkers that fuse histone dimers into single polypeptides, forming H4–H3 and H2B–H2A pairs. These connectors likely preserve the tetrameric and dimeric organization of the core in the absence of discrete subunits. Whether they also contribute to stability by constraining histone motion or reinforcing DNA wrapping, however, remains unresolved. If they reduce internal flexibility or support weakened interfaces, the connectors could act as structural compensators. If not, their role may be limited to preserving subunit continuity. We quantified structural variability by computing pairwise RMSDs for each connector across simulation frames for each connector type (Figure S8). The H2B–H2A connectors, located opposite the nucleosomal dyad, showed limited variability, with RMSDs predominantly below 4 Å. In contrast, the H4–H3 connectors displayed substantially greater variability, with values centered around 8–8.5 Å and extending up to 12 Å. This enhanced flexibility was consistent across simulations and most pronounced at the Exit side, where DNA unwrapping was more extensive. These results suggest that H4–H3 connectors exhibit more conformational heterogeneity than their H2B–H2A counterparts, potentially allowing them to respond more dynamically to changes in DNA–histone interactions during unwrapping.

To further investigate the the H4–H3 connector dynamics and how they relate to DNA unwrapping and histone motion, we clustered their conformations. Clustering analysis revealed that H4–H3 connectors adopt multiple structural states that reorganize extensively during DNA unwrapping. Fully wrapped DNA states were dominated by connector conformations with stable intrahistone hydrogen bonds involving loop regions of H3 and H4, and occasional interconnector contacts (Figure 6). As unwrapping progressed, these hydrogen bonding networks were gradually disrupted, accompanied by increased flexibility in connector positioning, including displacement of the H3 loop away from the histone core and rearrangement of stabilizing contacts. At higher unwrapping levels, hydrogen bonds near H3 reformed, suggesting local re-stabilization. Similar transitions were observed independently at both Entry and Exit sides, where distinct connector conformations emerged at each stage of unwrapping. Each was defined by unique hydrogen bonding patterns, consistent with the asymmetric DNA unwrapping established earlier. Detailed occupancy statistics and hydrogen bond profiles for each cluster are reported in the Supplementary Information (Section S1).

**Figure 6:**
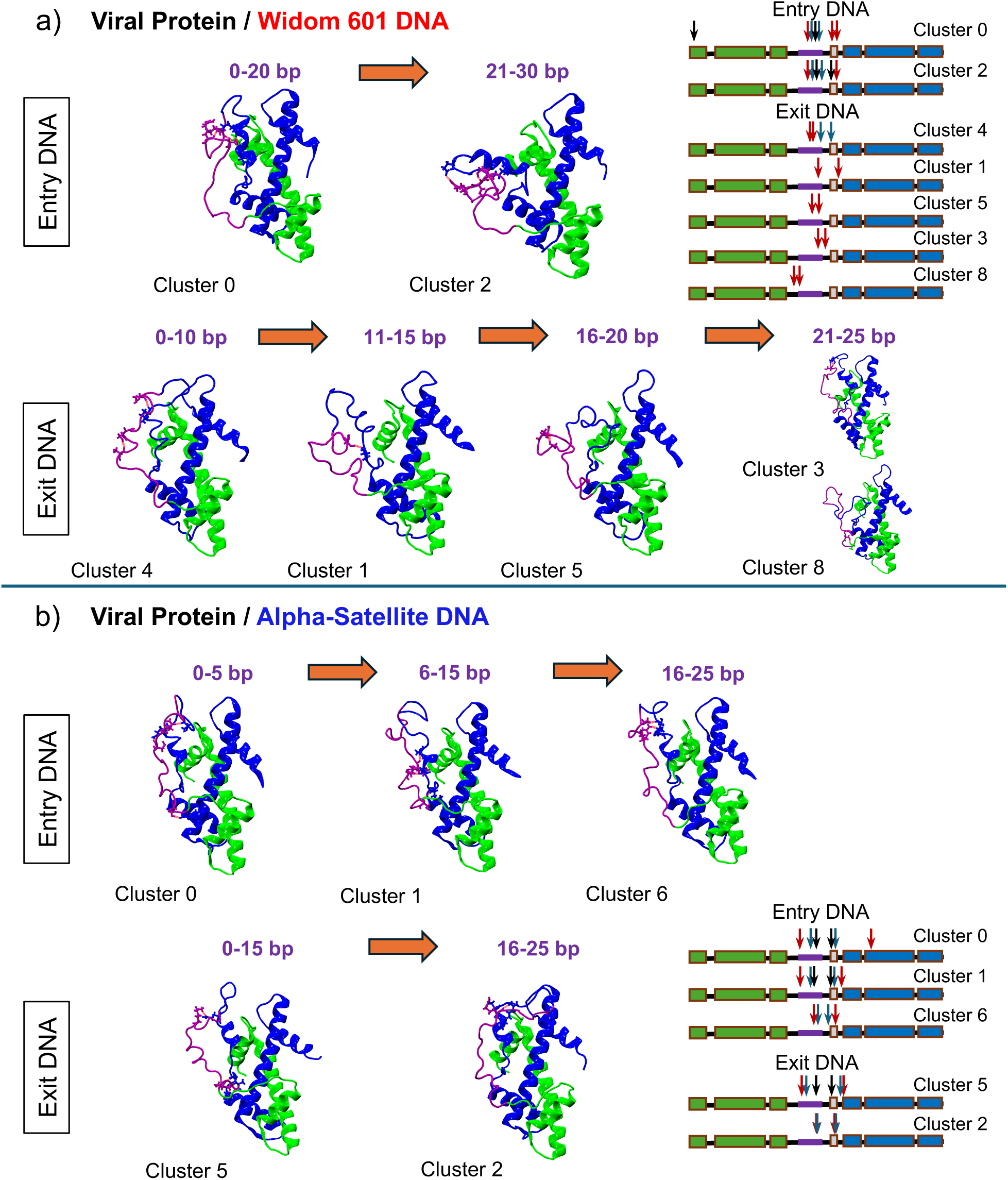
Cluster maps of fused H4–H3 histone connector conformations during DNA unwrapping. For both **(a)** Widom 601 and **(b)** Alpha-Satellite DNA, cluster maps show the dominant H4–H3 connector conformations at discrete Entry and Exit DNA unwrapping stages in viral systems. Arrows indicate transitions between stages. Representative structures for each cluster highlight key hydrogen bond positions within the fused histone interface, shown as arrows on accompanying structural block diagrams.

The extent of these conformational shifts depended on DNA sequence. Widom 601 nucleosomes sampled a broader range of connector states with more frequent hydrogen bond disruption, shifting, and reformation across unwrapping states, indicating greater flexibility, particularly between the connector and H3. Alpha-Satellite systems, however, featured more constrained conformations at both the Entry and Exit sides. Fully wrapped and early unwrapping states maintained an extensive hydrogen bond network, with both intrahistone and interconnector bonds formed along the length of the connector, primarily with H3’s loop and a-helix regions. As DNA unwrapped, these bonds progressively shifted to the top of the connector, largely involving intrahistone contacts with the H3 loop, while contacts at the bottom and interconnector bonds were lost. These trends indicate that viral nucleosome connectors flexibly respond to local structural changes during DNA unwrapping, and that the magnitude and pattern of this adaptation may be influenced by the underlying DNA sequence.

Finally, we assessed whether viral histone connectors provide thermodynamic compensation for the absence of canonical histone tails. MM/GBSA interaction energies were calculated between DNA and the viral connectors, and compared to those between DNA and individual histone tails in the full eukaryotic nucleosome (Table 3). In both cases, viral connectors formed relatively weak interactions with DNA. The H4–H3 connector, for example, had an energy of –21.6*±*11.2 kcal/mol, nearly tenfold lower than the H3 tail (–258.5*±*13.0 kcal/mol) and about sixfold lower than the H4 tail (–127.9*±*11.7 kcal/mol). A similar pattern was observed for the H2B–H2A connector, which also showed markedly weaker DNA interaction energies than its eukaryotic counterparts. These findings demonstrate that while viral connectors may offer modest stabilization, they fall far short of compensating for the loss of tail-mediated DNA interactions, reflecting a fundamental shift in the energetic architecture of viral chromatin.

**Table 3:**
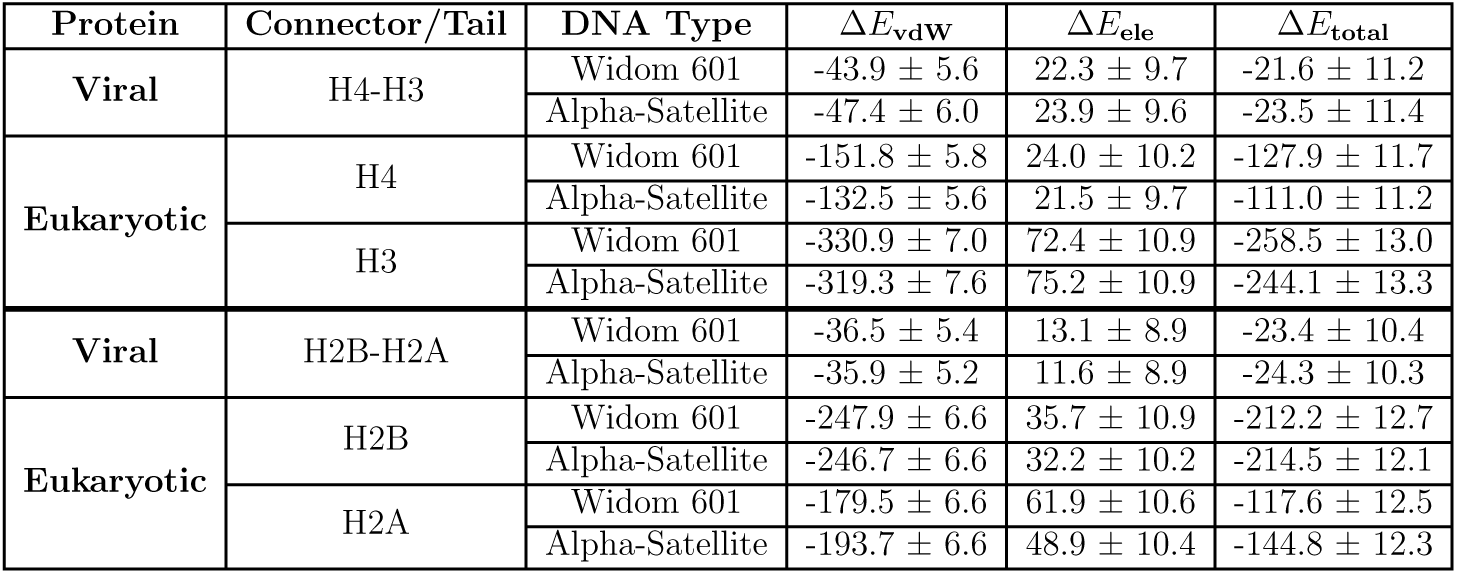
MM/GBSA Binding Energies (kcal/mol) — Connector/Tail–DNA Interface. Viral histone connectors exhibit significantly weaker DNA interaction energies compared to eukaryotic histone tails, indicating limited thermodynamic compensation for the absence of canonical tail–DNA contacts.

## Discussion

Viral nucleosome-like particles are structurally and thermodynamically less stable than their eukaryotic counterparts, driven by specific losses in molecular cohesion. While previous studies have resolved viral nucleosomes at high resolution, they leave open how these assemblies behave dynamically and how their physical features shape chromatin accessibility over time. Our simulations address this gap by showing that Melbournevirus nucleosomes have enhanced structural flexibility and reduced molecular cohesion, features that may reflect adaptive constraints on viral genome organization. Across all systems, we observed asymmetric and elevated DNA unwrapping, increased histone core fluctuations, and reduced protein compaction over time. These effects stem from weakened intra- and interhistone hydrogen bonding, diminished salt bridge formation at subunit interfaces, and a lack of stabilizing histone tails. Melbournevirus particles partially compensate through covalent peptide linkers that fuse histone dimers into single polypeptides. Although these connectors do not form extensive DNA contacts, they exhibit dynamic, state-dependent intramolecular hydrogen bonding. In tightly or extensively unwrapped states, the connectors adopt compact, hydrogen bond-rich conformations, suggesting they adjust to local structural strain.^30^

This structural responsiveness extends to the DNA itself. Viral nucleosomes exhibited frequent, asymmetric unwrapping of up to 20 base pairs on each DNA end, with Entry and Exit regions sampling both transient breathing-like states and more sustained unwrapped configurations. These transitions occurred stochastically and rarely rewrapped, resulting in broad distributions of end-to-end distance and radius of gyration. While Widom 601 and Alpha-Satellite DNA are strong positioning sequences^60^ in eukaryotic systems, they displayed distinct unwrapping behaviors in viral contexts, highlighting the sequence sensitivity of this chromatin architecture. Widom 601 favored unwrapping at the Entry and showed greater total unwrapping overall, while Alpha-Satellite unwrapped primarily from the Exit and maintained more compact structures. The enhanced breathing observed here reflects the looser histone-DNA coupling described above: even with favorable sequences, the reduced energetic drive for wrapping allows these systems to sample a wider range of conformations. The result is a dynamic ensemble of partially unwrapped states whose behavior is shaped by DNA sequence, local flexibility, and histone plasticity.

In contrast to the fluid interface dynamics seen in viral particles, eukaryotic nucleosomes maintain tighter control over DNA positioning^60^ through persistent histone-DNA interactions and extensive hydrogen bonding networks, many of which involve flexible histone tails.^22,23^ These tails contribute to overall compaction and modulate unwrapping dynamics by stabilizing wrapped configurations and limiting large-scale excursions. Even when tails are removed in eukaryotic systems, DNA unwrapping tends to be gradual and more symmetric, without the high-amplitude excursions or long-lived unwrapped states observed in viral particles. This suggests that the energetic landscape of viral nucleosomes is inherently less biased toward the wrapped state, leading to greater sensitivity to local sequence features and increased structural heterogeneity.

These findings highlight critical avenues for uncovering how viral chromatin reconciles instability with functional genome organization. First, while our study focuses on the most well-characterized viral nucleosome-like particle from Melbournevirus, it represents just one example within a broader evolutionary landscape. Over 1,500 viral histone proteins have been identified to date, forming primary structures ranging from histone singlets to quadruplets, each likely exhibiting distinct biophysical and dynamical modes of DNA compaction.^13,61,62^ Another key direction involves examining the effects of DNA methylation, which is widespread in large DNA viruses^63^ but was absent in our current models. Introducing methylated bases could test whether viral systems evolved to selectively stabilize modified DNA sequences, potentially revealing a compensatory role for DNA modifications in packaging. Likewise, mutational analysis of connector regions may clarify whether specific residues regulate their conformational flexibility or hydrogen bonding behavior during unwrapping transitions. While individual viral nucleosomes are inherently unstable, recent cryo-EM studies have shown that they form ultra-dense, tail-less arrays in the virion, wrapping approximately 121 base pairs per particle without linker DNA.^15^ This suggests that local plasticity may be an evolutionary adaptation that facilitates high-density chromatin packing,^64^ enabling tight genome compaction in the virion while retaining the ability to regain flexibility under host-like conditions. Coarse-grained modeling or simulations under varied ionic strengths could test this directly, revealing how viral nucleosomes shift between suppressed and permissive states. These directions will help clarify how dynamic, unstable particles can still support regulated genome organization in large DNA viruses.

Beyond their mechanistic differences, viral nucleosomes reveal distinct strategies for balancing genome compaction and accessibility. They are neither minimalist like archaeal chromatin nor canonical like eukaryotic nucleosomes, but instead define a third architectural mode. Archaeal histones form tail-less, polymeric structures without defined particles,^7^ while viral systems create compact nucleosome-like units stabilized by fused histone cores and structured connectors. By dissecting how Melbournevirus histones maintain compaction without terminal tails or canonical subunit interfaces, our work demonstrates the adaptability of the histone–DNA complex. Rather than representing a stripped-down core, viral nucleosomes show how chromatin features can be rewired to support both stability and dynamic access. Together, archaeal, eukaryotic, and viral chromatin illustrate the diversity of structural solutions to genome packaging and regulation. These systems provide a comparative framework for identifying which features are essential, which are flexible, and how physical constraints have shaped chromatin solutions across the tree of life.

## Acknowledgments

This project was supported by the National Institutes of Health grant R35GM119647.

## Author contributions

M.M and J.W. designed the project and developed the methodology. M.M. performed the molecular dynamics simulations, conducted the analyses, and prepared the visualizations. M.M. and J.W. co-wrote the original draft. J.W. supervised the project and secured funding.

## Data availability

Simulation input files, analysis scripts, and processed data are available at https://github.com/WereszczynskiGroup/supplemental-data-melo-viral-2025. Trajectory files have been deposited on Zenodo at https://doi.org/10.5281/zenodo.16342233. Waters have been removed and frames are temporally strided to reduce file size. These datasets are sufficient to reproduce all analyses and key results presented in the manuscript.

