## Supporting Information for "Viral Nucleosome-Like Particles Exhibit Dynamic Flexibility and Reduced Thermodynamic Stability"

### S1. Details of H4-H3 clustering analysis

Clustering was performed on H4–H3 connector side-chain conformations from viral nucleosomes assembled with Widom 601 and Alpha-Satellite DNA, using DBSCAN ($\varepsilon$ = 3.8, minPts = 4) to identify dominant structural states. After filtering out noise frames, 56,448 Widom and 65,156 Alpha-Satellite conformations were retained from an initial 160,000 simulation frames (80,000 from each DNA Entry and Exit site). Widom 601 nucleosomes displayed greater conformational diversity, with 21 total clusters and the eight most populated accounting for 33.3% of the data. In contrast, Alpha-Satellite systems showed more constrained sampling, with 17 total clusters and the eight most populated comprising 38.3% of frames (Table S2). These findings indicate that the viral connector is structurally responsive to DNA unwrapping and that this flexibility is modulated by DNA sequence.

DBSCAN cluster analysis further revealed that the relationship between DNA unwrapping and connector hydrogen bonding is both directionally asymmetric and sequence-dependent. In the viral Widom nucleosome (Figure 6a and Table S3), the Entry DNA side retained multiple native intrahistone hydrogen bonds up to 20 bp unwrapping. These included contacts within the connector itself, between the connector and the H3 $\alpha$N helix, and between the connector and the H4 $\alpha$1 helix, maintaining a compact conformation. Beyond 21 bp, the H4 contact was lost, and hydrogen bonding shifted primarily to the H3 $\alpha$N helix, with the connector-H3 unit moving away from the H4 region, consistent with a more open core conformation. On the Exit side, hydrogen bonding was more dynamic. During the 0-10 bp stage, only one intra-connector bond and one bond with the H3 loop were observed. At later stages, dominant clusters displayed stage-specific bonding: 11-15 bp involved H3 loop contacts, 16-20 bp shifted to intra-connector bonds, and 21-25 bp showed two distinct conformations, one bonded to the H3 loop near the top of the histone and one to the H4 region near the bottom. These transitions reflect a high degree of connector flexibility and asymmetry between Entry and Exit unwrapping.

In contrast, Alpha-Satellite nucleosomes (Figure 6b and Table S4) showed fewer dominant connector conformations and more persistent hydrogen bonding across all unwrapping stages. At least two bonds were maintained per unwrapping interval. At the Entry site, three consistent conformations were observed. Hydrogen bonds between the connector and H3 αN helix shifted slightly during unwrapping, with only minor repositioning of the lower connector region. At the 16-25 bp stage, one bond was lost, but the top of the connector remained stably associated with the H3 loop. The Exit side showed a single dominant conformation across both 0-15 bp and 16-20 bp stages, with hydrogen bonding transitioning from the H3 α1 helix to the H3 loop, mirroring Entry-side behavior. Overall, Alpha-Satellite DNA constrained connector motion and stabilized intrahistone interactions across unwrapping, in contrast to the greater flexibility and stage-dependent transitions observed with Widom DNA.

### S2. Supplemental Tables and Figures

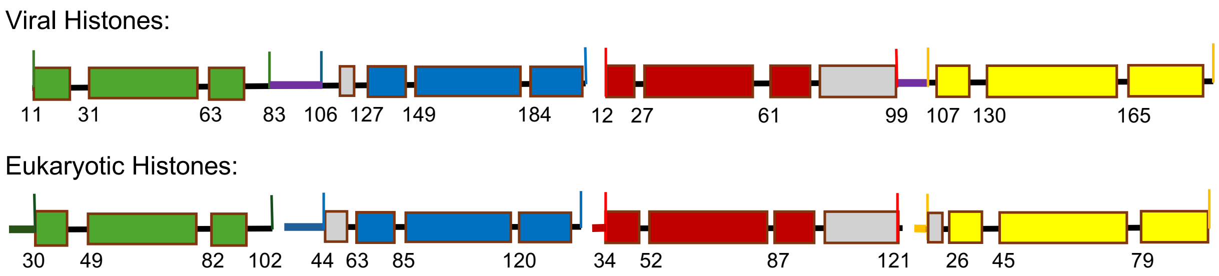


**Figure S1: Sequence alignment of core regions from viral and eukaryotic histones.** Thin vertical lines mark aligned residues across homologous segments of each histone fold domain. Thick horizontal bars indicate the positions of histone connectors in viral proteins or canonical tails in eukaryotic histones. Color scheme: H4 (green), H3 (blue), H2A (yellow), H2B (red).

**Table S1: MM/GBSA binding free energies between protein core–DNA interface (kcal/mol).** Viral protein cores show significantly weaker electrostatic and van der Waals interactions with DNA compared to eukaryotic nucleosomes. Differences between eukaryotic systems reflect the presence or absence of histone tails.

| Protein | DNA | $\Delta E_{\mathrm{vdW}}$ | $\Delta E_{\mathrm{ele}}$ | $\Delta E_{\mathrm{total}}$ |
|---|---|---|---|---|
| **Viral** | Widom 601 | -487.9 ± 11.1 | 90.7 ± 20.3 | -397.1 ± 23.1 |
| | Alpha-Satellite | -463.1 ± 10.9 | 94.1 ± 21.3 | -369.0 ± 23.9 |
| **Truncated Eukaryotic** | Widom 601 | -800.8 ± 9.8 | -125.3 ± 19.1 | -926.2 ± 21.4 |
| | Alpha-Satellite | -894.8 ± 8.9 | -132.3 ± 18.4 | -1027.2 ± 20.4 |
| **Eukaryotic** | Widom 601 | -829.6 ± 9.0 | -96.3 ± 18.0 | -925.8 ± 20.1 |
| | Alpha-Satellite | -865.7 ± 9.2 | -92.1 ± 18.3 | -957.9 ± 20.5 |

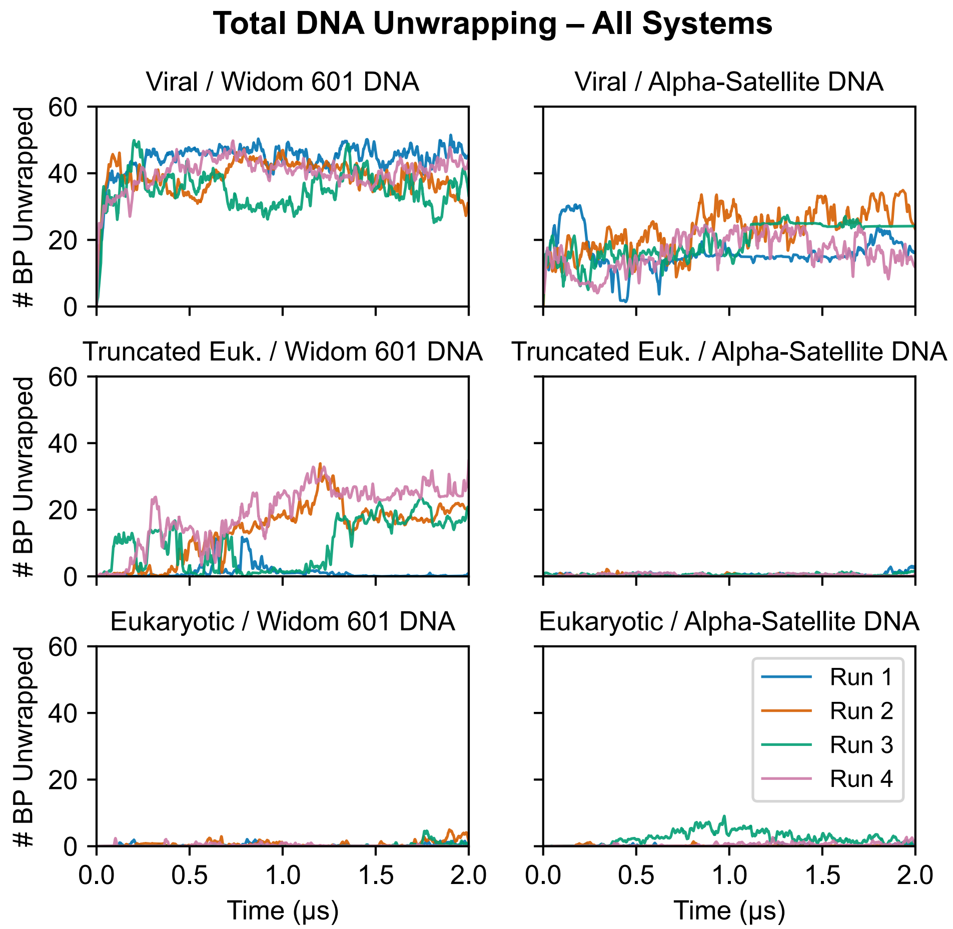


**Figure S2: Total DNA unwrapping in viral and eukaryotic nucleosomes.** Viral nucleosomes exhibit greater DNA unwrapping dynamics than their eukaryotic counterparts, with unwrapping peaking at ∼50 base pairs for the Widom 601 sequence and ∼35 base pairs for Alpha-Satellite DNA. In contrast, truncated eukaryotic nucleosomes reached a maximum of only ∼33 base pairs on the Widom sequence.

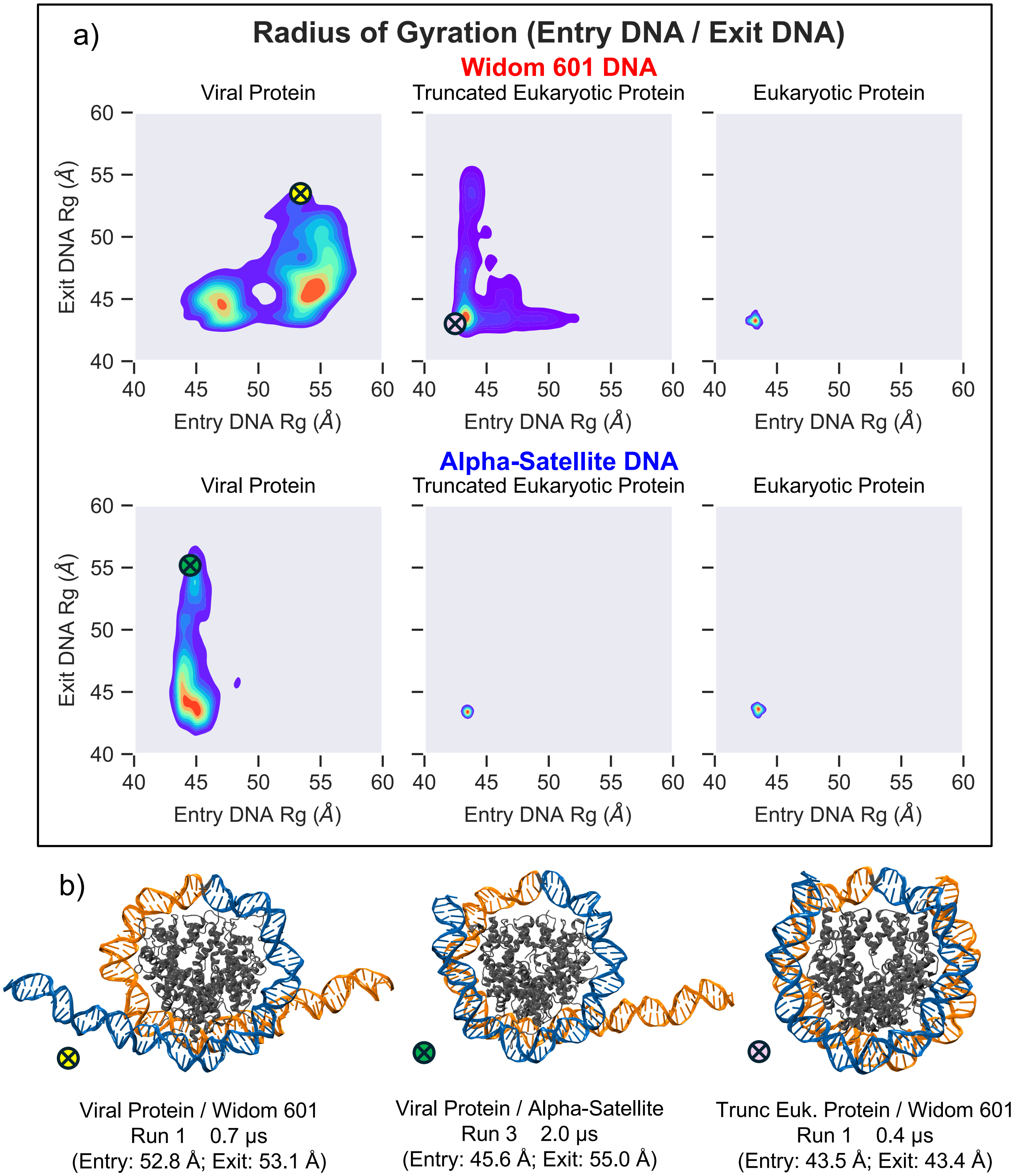


**Figure S3: Radius of gyration of nucleosome Entry vs. Exit DNA.** **(a)** The radius of gyration (Rg) of DNA relative to the dyad captures the spatial dynamics of Entry and Exit DNA ends, highlighting how DNA sequence and protein composition shape conformational variability. Viral nucleosomes exhibit higher Rg values, indicating greater DNA mobility than eukaryotic systems. **(b)** Representative snapshots depict Rg extremes.

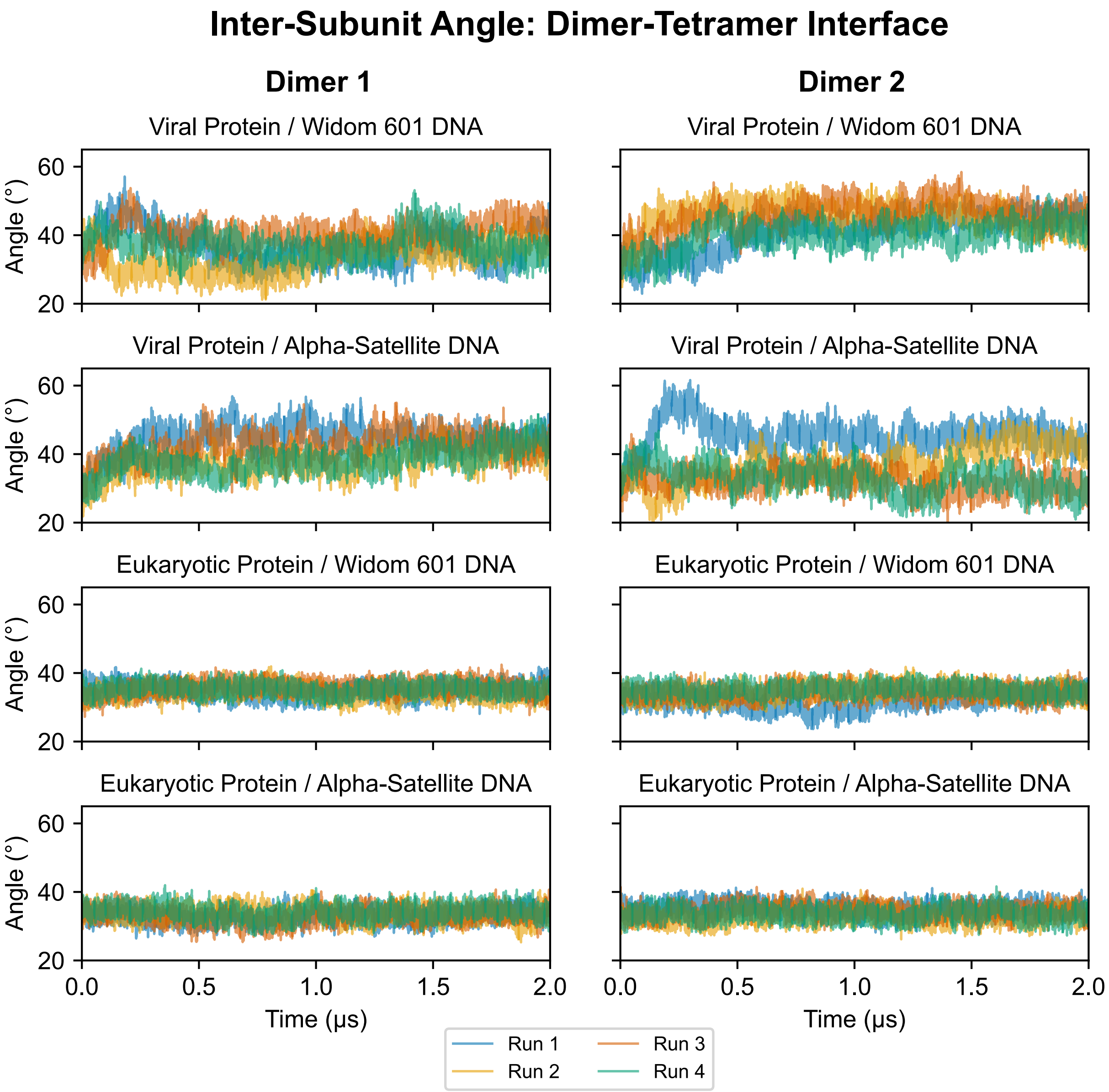


**Figure S4: Angular movement of dimers relative to tetramer in nucleosome core.** Comparison of dimer angular displacements relative to the H4–H3 tetramer reveals limited movement in eukaryotic nucleosomes, while viral systems exhibit increased flexibility with larger angular variations, reflecting a more dynamic histone core arrangement.

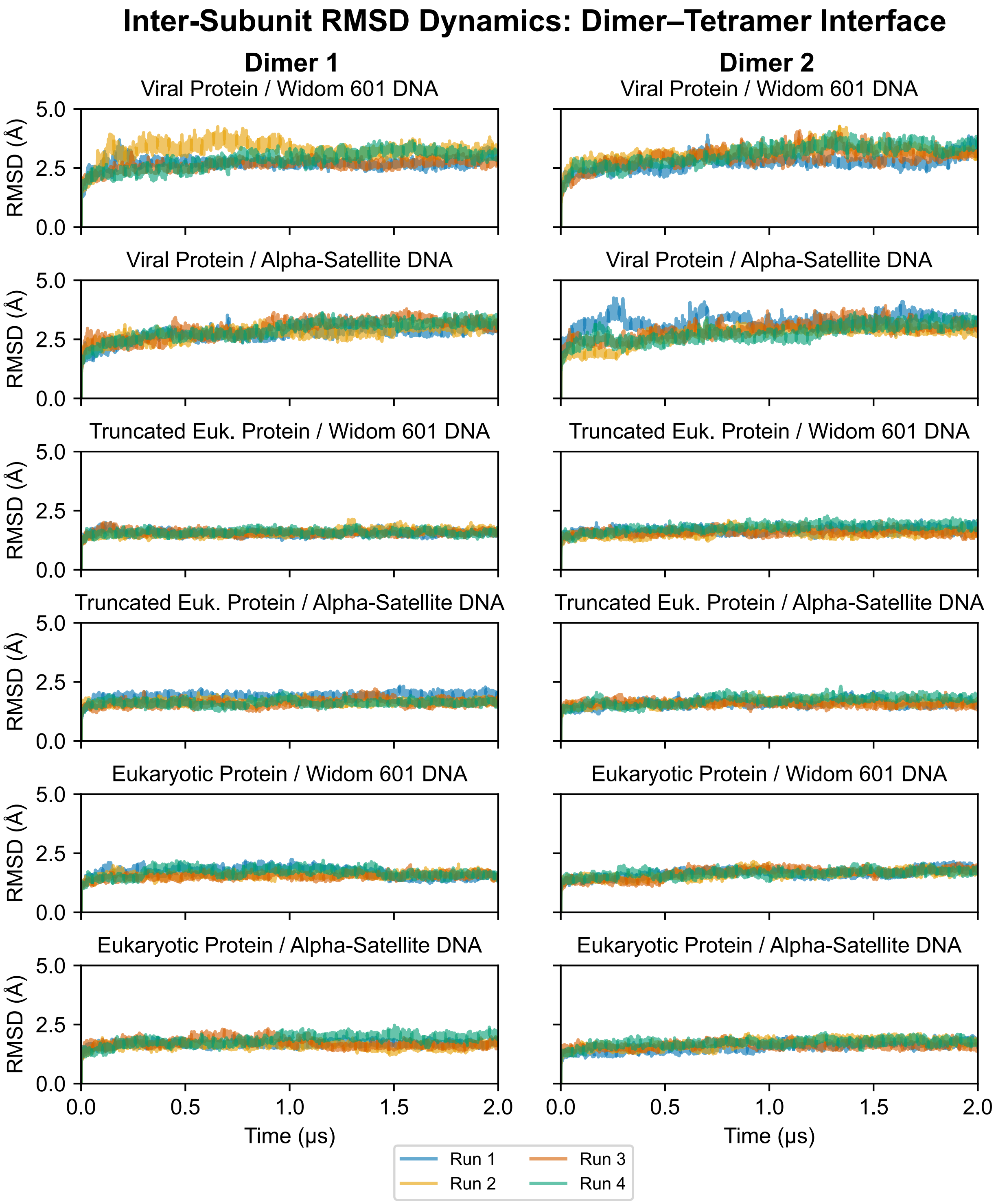


**Figure S5: RMSD of H2B–H2A dimers relative to H4–H3 tetramer across systems.** Root mean square deviations (RMSDs) of H2B–H2A dimers relative to the H4–H3 tetramer show minimal displacement in eukaryotic nucleosomes, whereas viral nucleosomes display broader RMSD distributions and greater dimer displacement, indicating enhanced structural heterogeneity.

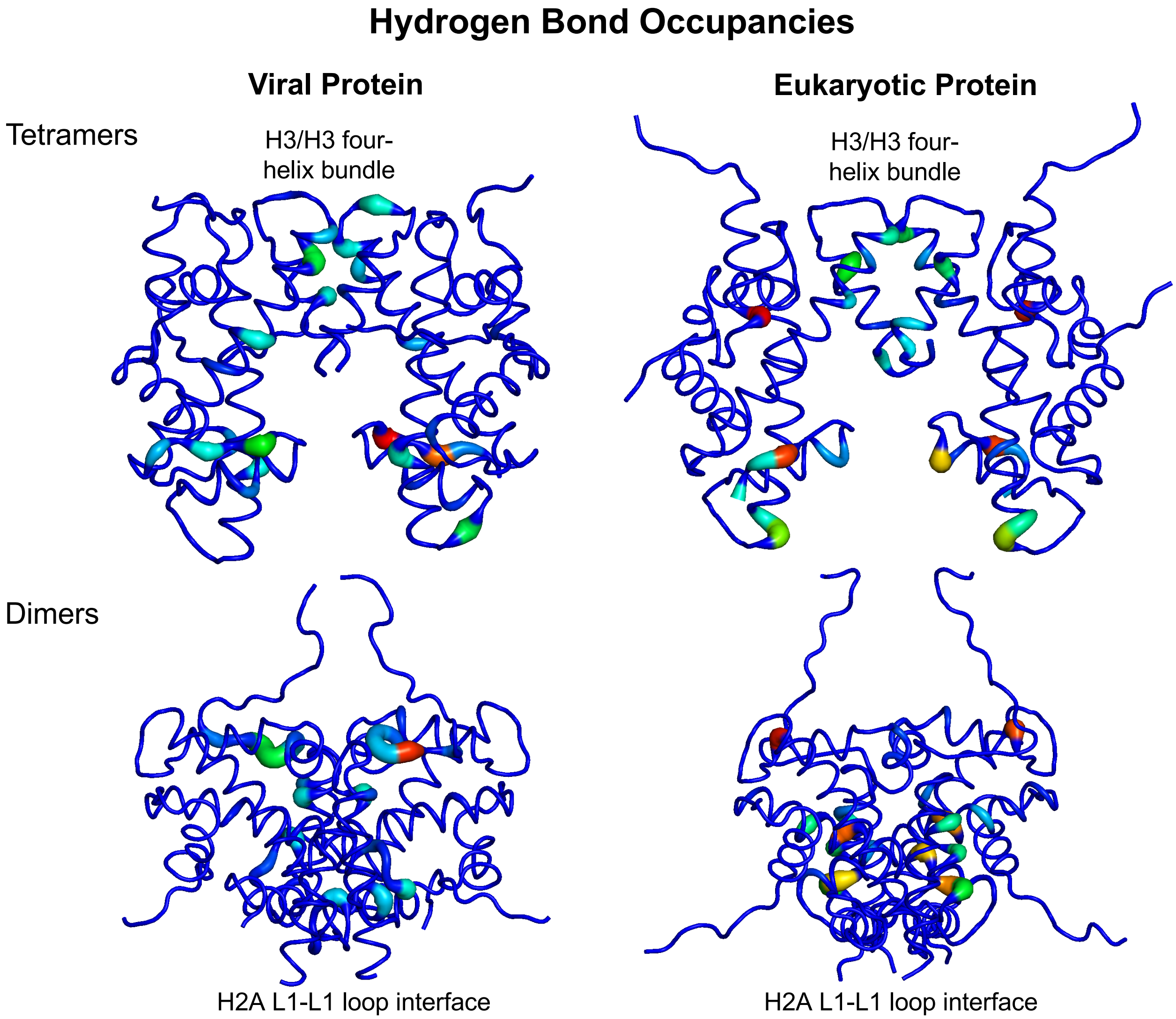


**Figure S6: Hydrogen bond occupancies in viral and eukaryotic proteins.** Structure-based models display residue-level hydrogen bond occupancies, visualized via B-factor scaling. The top row compares viral and eukaryotic tetramers; the bottom row shows viral and eukaryotic dimers. Key interaction sites, including the H3/H3 four-helix bundle and the H2A L1–L1 interface, are labeled.

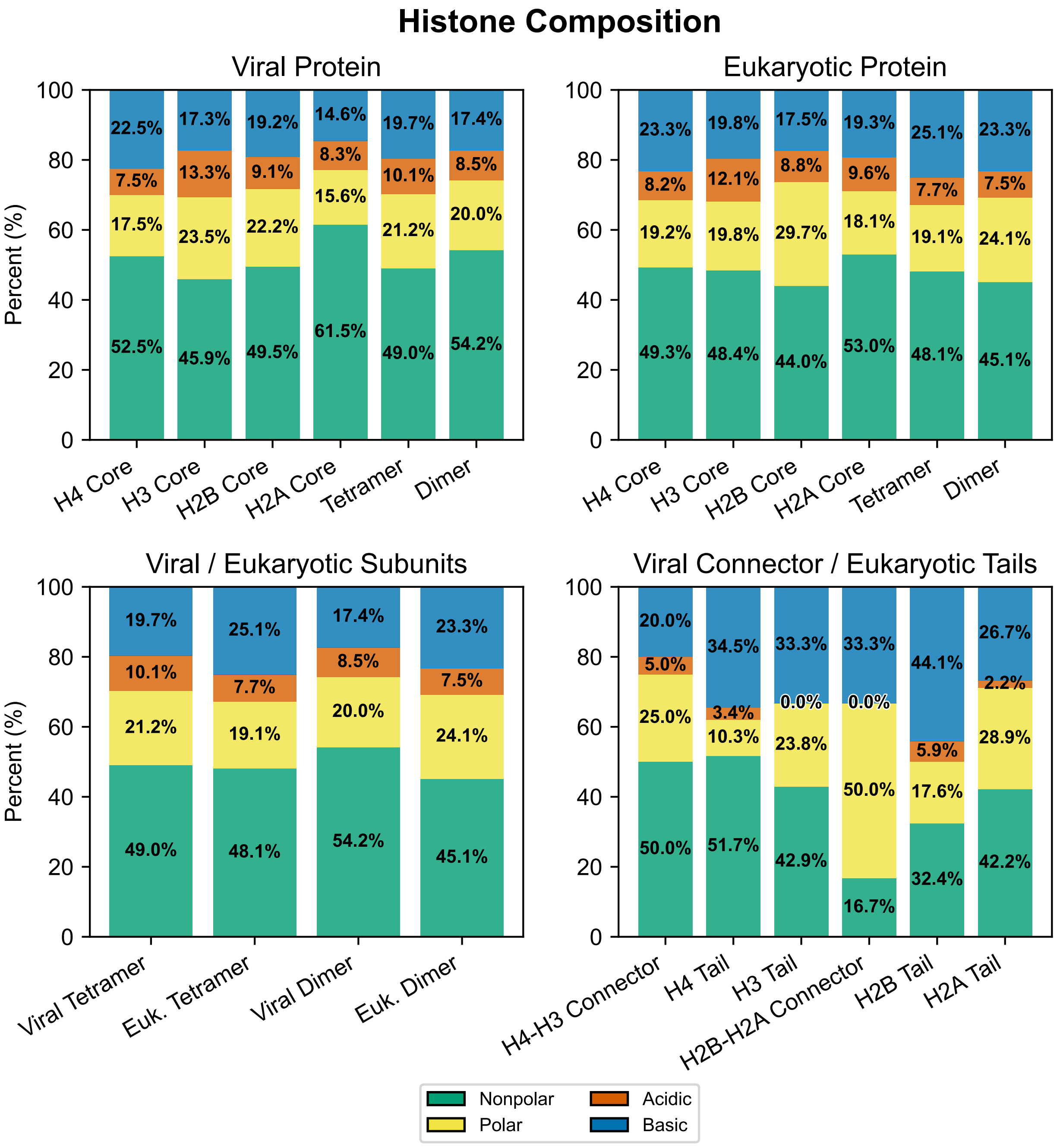


**Figure S7: Histone composition comparison between viral and eukaryotic proteins.** Viral and eukaryotic histones display similar amino acid compositions; however, differences in residue organization contribute to their divergent structural and dynamic properties.

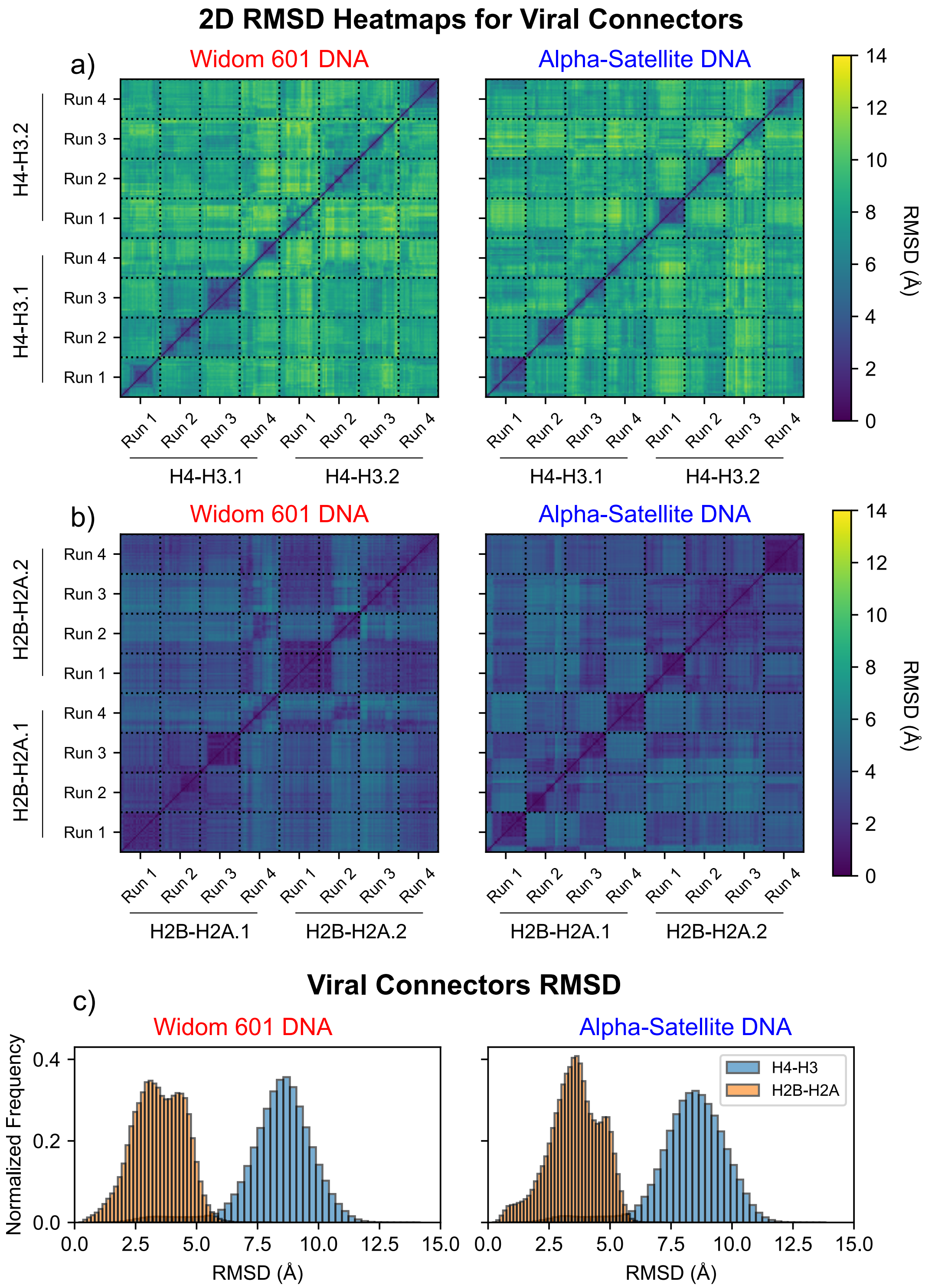


**Figure S8: Histone connector motion differences between viral H4–H3 and H2B–H2A.** **(a-b)** Two-dimensional RMSD heatmaps depict structural variability of viral **(a)** H4–H3 connectors and **(b)** H2B—H2A connectors for Widom 601 and Alpha-Satellite DNA systems. **(c)** RMSD distributions show that H4–H3 connectors exhibit substantially greater conformational variability (∼8 Å) compared to the more stable H2B-–H2A connectors (∼3 Å), reflecting enhanced flexibility of H4-–H3 during DNA unwrapping.

**Table S2: DBSCAN clustering of fused H4–H3 connector conformations in viral nucleosomes.** Clustering results of H4–H3 connector side-chain conformations from viral nucleosomes assembled with Widom 601 DNA (56,448 frames) and Alpha-Satellite DNA (65,156 frames), using DBSCAN with $\varepsilon = 3.8$ and minPts = 4. Frame counts represent non-noise frames retained from a total of 160,000 (80,000 Entry and 80,000 Exit frames).

| V-WIDOM | **Frames** | **Frac** | **AvgDist** | **Stdev** | **AvgCDist** |
|---|---|---|---|---|---|
| **Cluster0** | 16080 | 0.101 | 4.516 | 1.528 | 7.671 |
| **Cluster1** | 10612 | 0.066 | 4.964 | 1.629 | 8.340 |
| **Cluster2** | 7903 | 0.049 | 5.872 | 2.065 | 9.145 |
| **Cluster3** | 6489 | 0.041 | 7.740 | 1.710 | 6.769 |
| **Cluster4** | 4537 | 0.028 | 6.178 | 1.599 | 7.134 |
| **Cluster5** | 3621 | 0.023 | 5.229 | 1.916 | 8.031 |
| **Cluster6** | 1516 | 0.009 | 6.960 | 1.929 | 6.779 |
| **Cluster7** | 1282 | 0.008 | 3.655 | 0.865 | 8.722 |
| **Cluster8** | 1224 | 0.008 | 4.767 | 1.308 | 8.087 |

| V-ALPHA | **Frames** | **Frac** | **AvgDist** | **Stdev** | **AvgCDist** |
|---|---|---|---|---|---|
| **Cluster0** | 15485 | 0.097 | 4.065 | 1.313 | 7.124 |
| **Cluster1** | 12713 | 0.079 | 4.207 | 1.259 | 6.853 |
| **Cluster2** | 7682 | 0.048 | 3.981 | 1.330 | 6.911 |
| **Cluster3** | 6481 | 0.041 | 6.323 | 1.644 | 6.924 |
| **Cluster4** | 5547 | 0.035 | 5.099 | 1.432 | 7.248 |
| **Cluster5** | 5338 | 0.033 | 7.743 | 1.964 | 5.610 |
| **Cluster6** | 4229 | 0.026 | 5.421 | 1.306 | 6.254 |
| **Cluster7** | 2470 | 0.015 | 6.878 | 1.732 | 7.994 |
| **Cluster8** | 1445 | 0.009 | 6.587 | 1.470 | 5.783 |

**Table S3: Cluster populations across DNA unwrapping stages in viral nucleosomes with Widom 601 DNA.** Percentages of non-noise frames at each unwrapping stage assigned to the dominant fused H4–H3 connector clusters, shown separately for Entry and Exit DNA arms.

| V-WIDOM-ENTRY | **0-5bp** | **6-10bp** | **11-15bp** | **16-20bp** | **21-25bp** | **26-30bp** |
|---|---|---|---|---|---|---|
| **Cluster0** | **13.35%** | **56.54%** | **73.35%** | **41.42%** | 2.29% | 0.14% |
| **Cluster1** | 0.00% | 0.00% | 0.00% | 0.00% | 0.00% | 0.00% |
| **Cluster2** | 0.00% | 0.00% | 0.00% | 0.00% | **14.26%** | **10.40%** |
| **Cluster3** | 5.80% | 2.48% | 0.97% | 2.80% | 6.97% | 0.00% |
| **Cluster4** | 0.00% | 0.00% | 0.00% | 0.00% | 0.00% | 0.00% |
| **Cluster5** | 0.00% | 0.00% | 0.00% | 0.00% | 0.00% | 0.00% |
| **Cluster6** | 0.00% | 0.00% | 0.00% | 0.00% | 0.00% | 0.00% |
| **Cluster7** | 0.00% | 0.00% | 0.00% | 0.00% | 0.00% | 0.00% |
| **Cluster8** | 0.00% | 0.00% | 0.00% | 0.00% | 0.00% | 0.00% |

| V-WIDOM-EXIT | **0-5bp** | **6-10bp** | **11-15bp** | **16-20bp** | **21-25bp** |
|---|---|---|---|---|---|
| **Cluster0** | 0.00% | 0.00% | 0.00% | 0.00% | 0.00% |
| **Cluster1** | 0.42% | 3.53% | **28.21%** | 7.63% | 0.83% |
| **Cluster2** | 0.00% | 0.00% | 0.00% | 0.00% | 0.00% |
| **Cluster3** | 0.61% | 1.05% | 2.19% | 5.17% | **7.48%** |
| **Cluster4** | **7.88%** | **14.41%** | 5.68% | 0.01% | 0.00% |
| **Cluster5** | 0.05% | 0.29% | 1.78% | **18.77%** | 3.92% |
| **Cluster6** | 5.60% | 2.47% | 1.18% | 0.00% | 0.00% |
| **Cluster7** | 0.83% | 8.07% | 1.03% | 0.00% | 0.00% |
| **Cluster8** | 0.00% | 0.00% | 0.02% | 4.47% | **7.62%** |

**Table S4: Cluster populations across DNA unwrapping stages in viral nucleosomes with Alpha-Satellite DNA.** Percentages of non-noise frames at each unwrapping stage assigned to the dominant fused H4–H3 connector clusters, shown separately for Entry and Exit DNA arms.

| V-ALPHA-ENTRY | **0-5bp** | **6-10bp** | **11-15bp** | **16-20bp** | **21-25bp** |
|---|---|---|---|---|---|
| **Cluster0** | **33.82%** | 0.08% | 0.08% | 0.00% | 0.00% |
| **Cluster1** | 9.23% | **29.50%** | **22.67%** | 0.00% | 0.00% |
| **Cluster2** | 0.00% | 0.00% | 0.00% | 0.00% | 0.00% |
| **Cluster3** | 13.26% | 2.81% | 0.17% | 0.00% | 0.00% |
| **Cluster4** | 10.13% | 4.57% | 1.47% | 0.00% | 0.00% |
| **Cluster5** | 0.00% | 0.00% | 0.00% | 0.00% | 0.00% |
| **Cluster6** | 1.54% | 17.15% | 5.19% | **15.49%** | **20.00%** |

| V-ALPHA-EXIT | **0-5bp** | **6-10bp** | **11-15bp** | **16-20bp** | **21-25bp** |
|---|---|---|---|---|---|
| **Cluster0** | 0.00% | 0.00% | 0.00% | 0.00% | 0.00% |
| **Cluster1** | 0.00% | 0.00% | 0.00% | 0.00% | 0.00% |
| **Cluster2** | 0.00% | 0.00% | 1.78% | **24.33%** | **32.61%** |
| **Cluster3** | 0.00% | 0.00% | 0.00% | 0.00% | 0.00% |
| **Cluster4** | 0.00% | 0.00% | 0.00% | 0.00% | 0.00% |
| **Cluster5** | **10.18%** | **9.23%** | **8.15%** | 5.08% | 0.52% |
| **Cluster6** | 0.03% | 0.00% | 0.00% | 0.00% | 0.00% |
